# Short-term effects on brain functional network caused by focused-attention meditation revealed by Tucker3 clustering on graph theoretical metrics

**DOI:** 10.1101/765693

**Authors:** Takuma Miyoshi, Kensuke Tanioka, Shoko Yamamoto, Hiroshi Yadohisa, Tomoyuki Hiroyasu, Satoru Hiwa

## Abstract

This study examines the short-term effects of focused-attention meditation on functional brain state in novice meditators. There are a number of feature metrics for functional brain states, such as functional connectivity, graph theoretical metrics, and amplitude of low frequency fluctuation (ALFF). It is necessary to choose appropriate metrics and also to specify the region of interests (ROIs) from a number of brain regions. Here, we use a Tucker3 clustering method, which simultaneously selects the feature vectors (graph theoretical metrics and fractional ALFF) and the ROIs that can discriminate between resting and meditative states based on the characteristics of the given data. In this study, breath-counting meditation, one of the most popular forms of focused-attention meditation, was used and brain activities during resting and meditation states were measured by functional magnetic resonance imaging. The results indicated that the clustering coefficients of eight brain regions tended to increase through the meditation. Our results reveal that short-term effects of breath-counting meditation can be explained by network density changes in these eight brain regions.

## 1 Introduction

Mindfulness meditation is said to be influential in physicality, cognition, and mentality, and also has positive effects on well-being [1,2]. Mindfulness-based stress reduction (MBSR) [3] and mindfulness-based cognitive therapy (MBCT) [4] are clinical interventions based on mindfulness meditation, and it has been reported that these interventions alleviate symptoms of disorders, such as anxiety disorders [5,6], depression [7], and substance use disorders [8]. Furthermore, it has also been reported that the practice of meditation contributes to improvement of well-being, quality of life [1,2], immune function [9,10], and cognitive function [11–14], not only for unhealthy people but also for healthy ones.

On the other hand, the biological mechanisms of mindfulness meditation have also been studied. In particular, the neural basis of mindfulness meditation has been investigated using noninvasive neuroimaging methods such as electroencephalography (EEG) [15,16] and functional magnetic resonance imaging (fMRI) [17,18].

For example, Short et al. have reported that expert meditators showed higher activation in the anterior cingulate cortex (ACC) and dorsolateral prefrontal cortex (dlPFC) during meditation [19], indicating that long-term practice of meditation affects brain function through effects on neuroplasticity. Hasenkamp et al. have revealed that there were four cognitive states during meditation: FOCUS (representing maintenance of attentional focus on the breath), MW (representing mind wandering or loss of focus), AWARE (representing the awareness of mind wandering), and SHIFT (representing shifting of focus back to the breath). They also found that the different brain regions were activated in each cognitive state [20]. For example, in the MW state, the activity of the default mode network (DMN) [21,22], involved in self-related cognitive processing, was enhanced, while the activity of the salience network (SN) [23, 24], involved in current processing and detection of related stimuli, was strengthened in AWARE state. In FOCUS and SHIFT states, the activity of the central executive network (CEN) [24,25] (or the fronto-parietal attention network), involved in controlling attention resources, was enhanced. Their proposed model has been widely used to interpret the dynamics of the cognitive states during meditation. Furthermore, they have also reported that the functional connectivity between the dlPFC in the CEN and the right insula in the SN was higher in experienced, long-term meditators compared with short-term meditators [26].

From these previous studies, here we assume that there are two kinds of functional changes in the brain caused by meditation: 1) the persistent changes caused by long-term meditation practice (long-term effects), and 2) the non-persistent changes occurring temporarily for only short periods during meditation (short-term effects). Most previous studies have focused on expert meditators and studied these two effects by comparing them with novices. Moreover, most of the studies focusing on novice meditators have studied long-term effects by measuring brain activities of participants simultaneously with clinical intervention such as MBSR and MBCT [27,28]. However, the short-term effects of meditation on brain function in novice meditators have not been sufficiently studied. This is because meditating in a correct manner is not easy for novices and it is believed that repeated practice for at least several weeks is necessary to achieve a proper meditative state.

Here, we investigate the short-term effects of meditation on brain function in novice meditators. Mindfulness meditation mainly includes focused attention (FA) meditation, which keeps attention to specific objects, and open monitoring (OM) meditation, which monitors current experience without value judgments. Practicing FA meditation is easier than OM meditation, and sustaining attention with an intention is one of the most important components of mindfulness, which is why FA meditation is the first method taught in many meditation training programs [29–31]. In addition, it has been reported that breath-counting meditation, which is a kind of FA meditation, improved psychological state and also reduced MW in novice meditators [32]. Therefore, FA meditation is one of the best methods for novices and is also used in this study.

In recent years, functional connectivity analysis, which quantifies temporal correlation between different brain regions, has been used to investigate brain function [33,34]. Furthermore, graph theoretical analysis has often used to characterize the properties of functional networks where each brain region is regarded as a node and a connection between brain regions is regarded as an edge in a mathematical graph. A variety of graph metrics have been applied to functional brain networks associated with different cognitive states and experimental conditions [35,36]. The functional brain network characterizing the meditative state, and its network property, has already been investigated in expert meditators [20]. Therefore, in this paper, the graph theoretical metrics of function networks are analyzed for novice meditators only, and the difference in network properties between resting and meditative states are compared each to investigate how the short-term practice of FA meditation affects the functional network. However, since there are a wide variety of graph theoretical metrics, such as degree centrality, betweenness centrality, and clustering coefficient, it is necessary to select the most suitable ones, such that the short-term effects on the functional brain network are appropriately represented.

In addition, not only indicators showing relationships between brain regions, such as functional connectivity, but also those representing local activity, such as brain activation, are important for investigating brain function. Meanwhile, MBSR also has a minimum of 1-5 minutes of meditation practice [37], and it is thought that sustained meditation practice at least for several minutes is necessary to achieve a stable meditative state. Therefore, in the experimental design, the duration of the task (meditation) blocks was more than several minutes to measure the brain activity during meditation practice, which was very problematic in the activation analysis because low frequency signal of 0.01 Hz or less is usually removed as noise in the typical method based on general linear model (GLM). Incidentally, in resting-state fMRI studies, the amplitude of low frequency fluctuations (ALFF) [38] that represents blood-oxygen-level-dependent (BOLD) signal power within the frequency band of interest (0.01-0.1 Hz) has been used as the indicator of local brain activity, and its correlation with the brain activation has also been reported in recent years [39]. Thus, in this study, in addition to the graph theoretical indicators, fractional ALFF (fALFF), which is a modified version of ALFF, was used as the local activity indicator.

Although definition of brain region depends on the brain parcellation method, it is important but difficult to determine the region of interests (ROIs), because there are about 100 or more brain regions to be analyzed (e.g., 116 regions defined by automated anatomical labeling). Therefore, in this study, Tucker3 clustering (T3Clus) [40, 41], which simultaneously selects the feature vectors (graph theoretical metrics and fALFF) and the ROIs that best discriminate between resting and meditative states based on the characteristics of the given data, was used. T3Clus is a clustering method based on three-way principal component analysis using the Tucker3 model [42] and classifies data while eliminating irrelevant features by reducing the dimension of data.

Here, we explain the outline of this study. First, the brain activities of the novice meditators were measured during a 5-min resting state and 5-min breath-counting meditation using functional magnetic resonance imaging (fMRI). Then, three graph theoretical metrics and fALFF, calculated for each brain region of all participants in both of resting and meditative states, were projected to 2D space by T3Clus, and the brain regions and feature indicators characterizing the difference between the two state were extracted. This difference was regarded as the short-term effects of FA meditation in the novices, and its characteristics were investigated based on the selected feature vectors and ROIs.

## 2 Materials and Methods

### 2.1 Overview of the proposed method

Here we propose method to extract the meaningful feature vectors to distinguish between two experimental conditions, resting and meditative states. First, BOLD time courses during the two conditions were measured for all subjects using fMRI. Second, three network feature vectors: degree centrality, betweenness centrality, and clustering coefficient [35], and one local activity measure, fALFF, representing the intensity of spontaneous brain activity [43], were chosen and calculated to quantify the brain states. Finally, T3Clus was applied to classify the two experimental conditions, simultaneously decomposing original feature space into low dimensional space to maximize the classification accuracy. Here, we used the supervised T3Clus method, where the experimental conditions are used as the class label.

### 2.2 Subjects

Twenty-nine healthy subjects (aged 22.9±2.3 years, 6 females, all right handed) participated in this experiment. Total hours of the meditation practice for each of them were less than 30 hours, and none of them experienced daily meditation training. All participants were informed about the experimental method and the risk and signed written informed consents. This study was carried out in accordance with the research ethics committee of Doshisha University, Kyoto, Japan (approval code: 15098).

### 2.3 Data acquisition

Whole-brain imaging data were acquired with a 1.5 T MR scanner (Echelon Vega, Hitachi, Ltd., Tokyo, Japan). Functional images were obtained using gradient echo-echo planer imaging (TR=2500 ms, TE=40 ms, flip angle=90°, FOV=240 mm, 5.0-mm thick slices, matrix size=64×64, number of slices=25). We also employed an Rf-spoiled steady state gradient echo (RSSG) sequence to obtain T1-weighted structural images (TR=9.8 ms, TE=4.0 ms, flip angle=8°, FOV=256 mm, 1.0-mm thick slices, matrix size=256×256, number of slices=192).

### 2.4 Experiment

The experiments consisted of a 5-min resting state block (pre-rest), a 5-min meditation block, and a 10-min resting state block (post-rest) as shown in Figure 1. Start and stop of meditation was informed by an auditory signal via headphones. The total duration of the experiment was 20 minutes. Participants practiced a simple-guided breath-counting meditation for a few minutes before entering the fMRI scanner.

**Figure 1:**
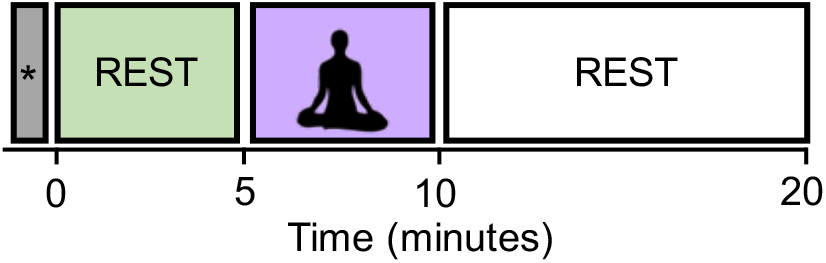
The experimental design. The block labeled with the symbol contains the dummy six volumes (with the duration of 15 s) acquired before the start of the first resting block. They were excluded from the analysis in order to eliminate the non-equilibrium effects of magnetization.

**Figure 2:**
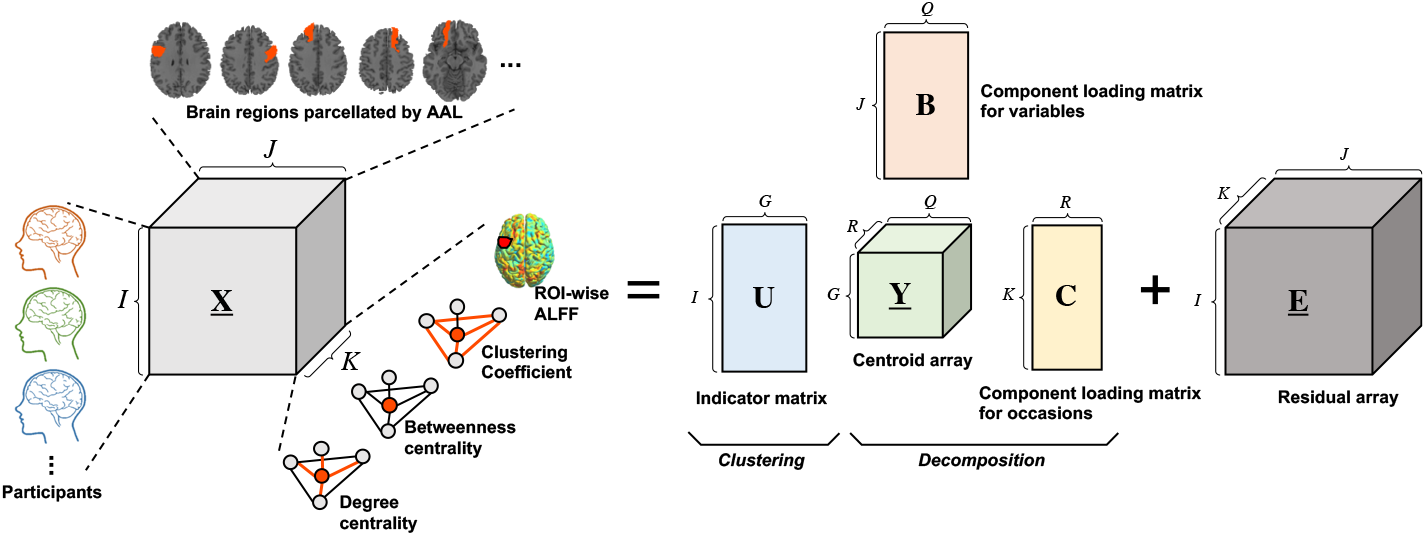
Schematic illustration of Tucker3 clustering on functional neuroimaging data. **<u>X</u>**(*I* × *J* × *K*), **<u>Y</u>**(*G* × *Q* × *R*) and **<u>E</u>**(*I* × *J* × *K*) denote the three-way data array of observation data, the centroid array, and the residual array, respectively. **U**(*I*×*G*), **B**(*J*×*Q*), and **C**(*K*×*R*) denote the indicator matrix defining a partition of the objects into *G* classes, the component loading matrix for variables, and the component loading matrix for occasions, respectively.

In the breath-counting meditation block, participants were asked to breathe through their nose and to try not to change the breath interval. They counted their breath silently from one to ten.. They were also instructed to restart counting from one if they got to ten, or if their mind got distracted. Their eyes were kept closed consistently in the scanner. In the rest block, they were instructed to stay relaxed without focusing on their breathing. The dummy six volumes were acquired before the start of the first resting block and also excluded from the analysis in order to eliminate the non-equilibrium effects of magnetization.

We focused on the pre-rest and meditation blocks to see the short-term effects of meditation. The post-rest block will be used to study another hypothesis, but was excluded for the current study.

### 2.5 Data preprocessing

The fMRI data were preprocessed using SPM12 (Wellcome Department of imaging Neuroscience, London, UK) on MATLAB (MathWorks, Sherborn, MA). All functional images were corrected for motion effects using a six-parameter rigid body linear transformation and slice-time corrected to the middle slice. Then, T1-weighted anatomical images were coregistered to the mean of the corrected functional images. These functional images were normalized to the Montreal Neurological Institute/International Consortium for Brain Mapping (MNI/ICBM) standard and spatially smoothed using an isotropic Gaussian filter (FWHM = 8 mm). To minimize global drift effects, the signal intensities in each volume were divided by the mean signal value for the run and multiplied by 100 to produce percent signal change from the run mean.

In addition to preprocessing through SPM12, the following, additional, pre-processing was performed. First, the whole brain was divided into gray matter, white matter, and cerebral spinal fluid (CSF). Then, nuisance regression was performed based on an anatomical component-based noise correction method (aCompCor) [44] to regress out the mean BOLD signals from the white matter and CSF and also remove head-movement and physiological confounding effects. Here, the main task effect from the meditation block (modeled as a canonical hemodynamic-response-function-convolved response) and its first order derivative were also regressed out to remove any potential confounding effects of shared task-related responses. Finally, a band-pass filter (0.008–0.09Hz) was applied to reduce the effects of physiological and low-frequency noise.

The preprocessed functional images were parcellated into 116 regions defined by automated anatomical labeling (AAL) and the mean BOLD time course was calculated for each region. These 116 time-courses were used to calculate the feature vectors.

### 2.6 Graph theoretical functional network analysis

Functional connectivity analysis is used to measure statistical interdependence (mutual information), without explicit reference to causal effects, by measuring correlation/covariance, spectral coherence, or phase-locking of time series brain activity between each brain region [34]. In this study, Pearson correlations were calculated between ROI-wise BOLD time courses during pre-resting and meditation blocks, and 116 × 116 functional connectivity matrices (FCMs) were obtained for each participant in each of the two conditions. The obtained FCMs were transformed to convert the sampling distribution of the Pearson correlation into a normal distribution. They were also binarized to determine the presence or absence of functional connections between the 116 regions. Determination of the appropriate thresholding method and its threshold value is one of the most crucial issues in graph theoretical functional network analysis. We used cost-based thresholding where the ratio of the connections with the strongest connectivity values out of the possible number of connections are preserved (e.g. cost 0.100 means the strongest 10% of possible of connections exist in the thresholded matrix). For threshold determination, we calculated the different threshold settings (from 0.050 to 0.500, increments: 0.025), and, then, the following three criteria were applied to them to choose the single threshold setting for the analysis.

#### 1) Small-world characteristics

Based on the assumption that small world topology are seen in the functional networks [45–47], we calculated the global and local efficiency measures for all threshold settings, and then found that the cost range: 0.050 - 0.425 satisfied the small-world network criteria: higher global efficiency than a lattice graph but lower global efficiency than a random graph.

#### 2) Similarity to the average characteristics

We aimed to determine the single threshold that can extract the network with group average characteristics between the different threshold settings (cost: 0.050 - 0.425). Therefore, we calculated the degree centrality measure for each brain region, for all participants, in each condition. Then, we averaged the degree centrality of each region within participants, which were used as across-subject average characteristics for the different cost thresholds. To determine the single threshold that was the most similar to the average characteristics, Pearson’s correlation between the across-subject average degree distribution and the degree distribution of the FCM, thresholded by a certain cost value, were calculated for each cost setting. The cost value with the highest correlation with the across-subject characteristics was chosen.

#### 3) Stability in the community structure

If the network structure are stable across the cost, the number of communities existing in the thresholded network should be also stable [48]. Therefore we used the number of communities detected by Newman clustering [49] on the thresholded network as a cost criterion. The community clustering was performed for all the participants, all cost settings, in each condition. The standard deviation of the number of communities across participants was calculated for each cost. The cost setting with the smallest deviation was chosen for the stable cost threshold.

According to these three criteria, we got reasonable setting: cost = 0.225, and we reported results for the cost dertermination in Supplementary Figure S1. Three graph theoretical metrics: degree centrality, betweenness centrality, and clustering coefficient were calculated as the indicators quantifying the network characteristics of each region [35].

### 2.7 Fractional ALFF calculation

We used fALFF measure to quantify the intensity of spontaneous brain activity induced or altered through meditation. ALFF is derived by calculating the sum of amplitudes of low-frequency of a specific frequency range, which is regarded as including spontaneous brain activity (i.e., 0.008-0.09 Hz was chosen for this study). It has been reported that the ALFF of the DMN is significantly higher in the resting state [38]. The fALFF, used in this paper, is the modified form of ALFF obtained by dividing ALFF by the sum of the amplitudes of all the possible frequency bands to reduce the influence of noise [43]. In this paper, fALFF were calculated for all voxels using CONN toolbox and then averaged within the brain region defined by AAL.

### 2.8 Tucker3 clustering for brain state classification

T3Clus is a the clustering method that simultaneously performs three-way principal component analysis and k-means clustering on the principal component scores [40, 42]. Three-way data means that multiple objects were observed foron multiple variables oin multiple occasions. Here, a set of observation variables, X, are is organized as *I* × *J* × *K* three-way array, where *I*, *J* and *K* denote the number of objects, variables, and occasions, respectively. With an increase of dimensionality of each of the three modes (objects, variables, occasions), the difficulty of the objects (data) classification will be increased [50]. In conventional approaches, one of the easiest ways to tackle this problem is to reduce the dimensionality by principal component analysis etc., and then perform the clustering. However, such an approach can result in loosing important prominent information which that contributes to classification through dimensionality reduction [51,52]. In theis case of, T3Clus, in which the dimensionality of the variables and occasions are reduced so that the objects are the most classified by combining three-way principal component analysis and k-means clustering are effective.

The Tucker3 model is represented by

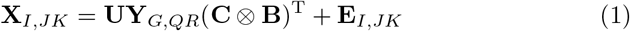

where **X**_*I,JK*_(*I*× *JK*), **Y**_*G,QR),G×QR*_, and **E**_*I,JK*_(*I× JK*) denote the ‘matricized’ versions of the **<u>X</u>**(*I* × *J* × *K*), the centroid array *<u>Y</u>*(*G* × *Q* × *R*), and the residual array **<u>E</u>**(*I* × *J* × *K*), respectively. **U**(*I* × *G*), **B**(*J* × *Q*), and **C**(*K* × *R*) denote an indicator matrix defining a partition of the objects into *G* classes, a component loading matrix for variables, and ca component loading matrix for occasions, respectively, and ⊗ denotes the Kronecker product of matrices. AdditionallyBesides, *G*, *Q*, and *R* denote number of classes, components for variables, and components for occasions, respectively.

T3Clus solves the following optimization problem:

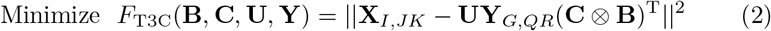

subject to **B** and **C** column-wise orthonormal and **U** binary and row-stochastic.

In this study, functional neuroimaging data were treated as three-way data consisting of 29 participants, 116 brain regions, and four regional functional metrics, degree centrality, betweenness centrality, clustering coefficient, and fALFF. Here, generally, indicator matrix U is constrained to have only one nonzero element per row to indicate which class each row belongs to, and it is optimized for solving the above optimization problem. However, in this study, we determined each element of U without the optimization process because it was obvious which experimental conditions (i.e., resting or meditative states) each data belonged to.

It should be noted that T3Clus leads to derivation of linear transformation from the original data space into low-dimensional space by optimizing the Kronecker product of *C* and *B*. Therefore, the component scores after executing T3Clus can be obtained by:

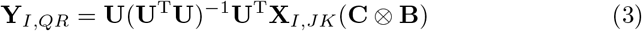

In this study, the 29 participants’ data were classified into two classes, simultaneously with reducing the number of brain regions from 116 to 2 and the number of functional metrics from 4 to 1 dimension using T3Clus (i.e., *G* =2, *Q* = 2, and *R* =1). As a result, the two-dimensional component scores of the neuroimaging data during the resting and meditative states of the 29 participants were obtained and were plotted in two-dimensional space.

### 2.9 Finding discriminating brain regions and their features

Since the component scores are obtained by equation (3), the Kronecker product **C** ⊗ **B** can be regarded as a weight vector forto the observation data. Thus, its elemental value can also be treated as the metric indicating thean importance of each row or column of the three-way array. In the meditation dataset, the higher elemental value indicatesrepresent that the corresponding feature is important forprominent to discriminating betweene the brain states between under resting- and meditation conditions.

Here we aim to elucidateobtain the essential combination of the brain regions and their functional brain metrics by analyzing the component loading matrix **C** ⊗ **B**. That is, discriminating features are extracted from the elements of the component-loading matrix whose values are necessary to discriminate between resting- and meditation conditions.

In order to achieve this, we appliedy the permutation tests [ [53–57] toon the component-loading matrix. The class labels of the state were randomly permuted 10000 times and T3Clus was applied toon the data set of each permuted label. The probability that loading with a larger absolute value than the loading obtained from original data set was given was the p-value. The significant brain regions and feature values were extracted under p-values < 0.05.

### 2.10 Performance comparison with the conventional approach

To ensure the effectiveness of applying T3Clus to our study in terms of finding the prominent features to discriminate different cognitive conditions, the paired t-test was applied between resting and meditative state in each of four feature values, and then the brain regions whose feature values were significantly different between two conditions were extracted (*p* < 0.05, FDR-corrected) and compared with the results obtained by T3Clus.

## 3 Results

### 3.1 Paired t-test

The paired t-test performed on each of the four feature metrics for each brain region between resting and meditative states revealed that there were no significant differences between two states in all brain regions for all feature values (*p* > 0.05).

### 3.2 Low-dimensional representation of the brain states derived by T3Clus

Figure 3 shows the two-dimensional representation of each participant’s brain state during the two experimental conditions, obtained by T3Clus. Each axis was determined by T3Clus so as to maximize the difference between two conditions, simultaneously reducing the dimensionality of the brain regions and feature metrics.

**Figure 3:**
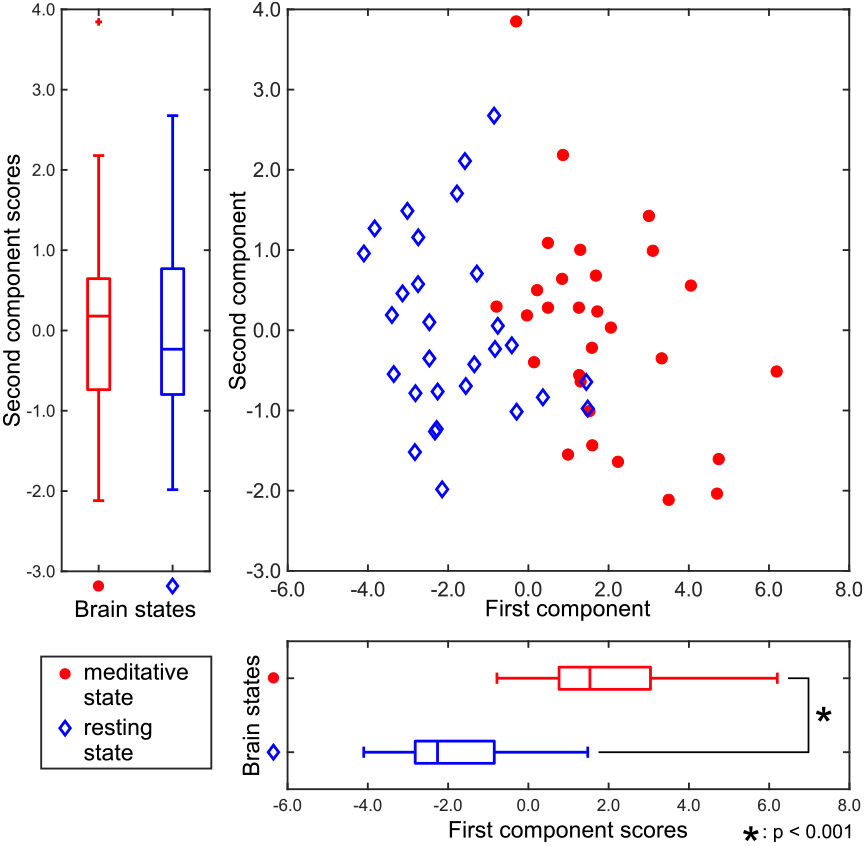
Component scores obtained by T3Clus. Each point indicates each participant’s data. The meditative state is red-colored and the resting state is colored blue.

There was a significant difference between the first component scores in the resting and meditative conditions (*p* < 0.001), while there was a no significant difference between the second component scores. Therefore it can be said that the discriminating brain regions and features to classify between two states exist in the component loadings for the first component. The component loadings for the first component are shown in Figure 4.

**Figure 4:**
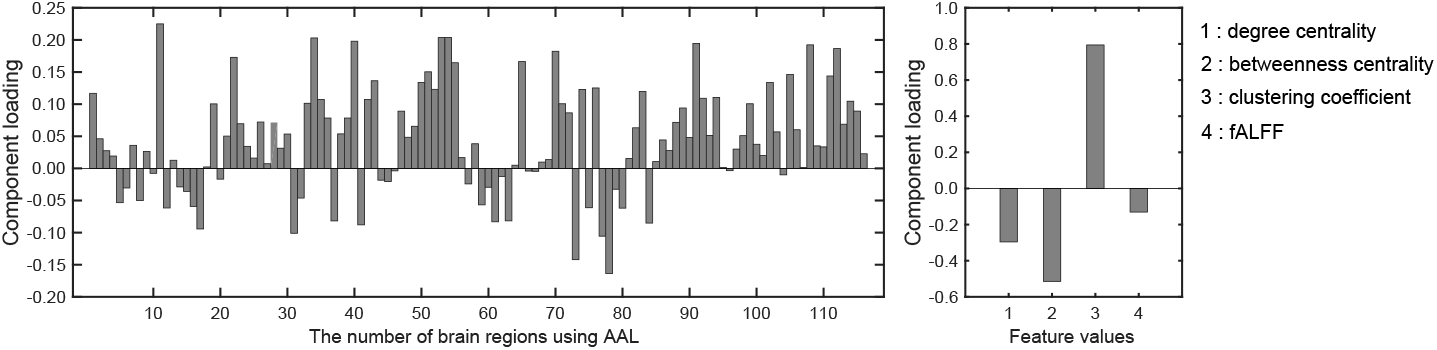
Component loadings obtained by T3Clus. Left: Loadings for brain regions. Right: Loadings for four feature values.

To extract the essential brain regions and their feature values, the permutation test was performed on the component loading matrix. Table 1 and 2indicate the significant brain regions and feature values with p-values < 0.05, respectively. The significant brain regions shown in Table 1 were the top 8 largest component-loading values in Figure 4. On the other hand, for the feature values, only the clustering coefficient was extracted as a significant discriminating feature.

**Table 1:** First component loading of brain regions (*p* < 0.05)

| Brain region | loading | p-score |
| --- | --- | --- |
| Frontal Inf Oper L | 0.2250 | 0.0125 |
| Occipital Inf R | 0.2036 | 0.0215 |
| ParaHippocampal R | 0.1981 | 0.0224 |
| Cerebelum 10 R | 0.1923 | 0.0281 |
| Cingulum Mid R | 0.2031 | 0.0284 |
| Cerebelum Crus1 L | 0.1946 | 0.0309 |
| Occipital Inf L | 0.2036 | 0.0324 |
| Paracentral Lobule R | 0.1824 | 0.0397 |

Therefore, these results indicated that the eight brain regions and their clustering coefficients were essential for discriminating brain states between resting and meditative conditions. Figure 5 shows the component loading maps of the eight brain regions that contribute to the classification of the two states, displayed on axial slices at different *z* levels.

**Figure 5:**
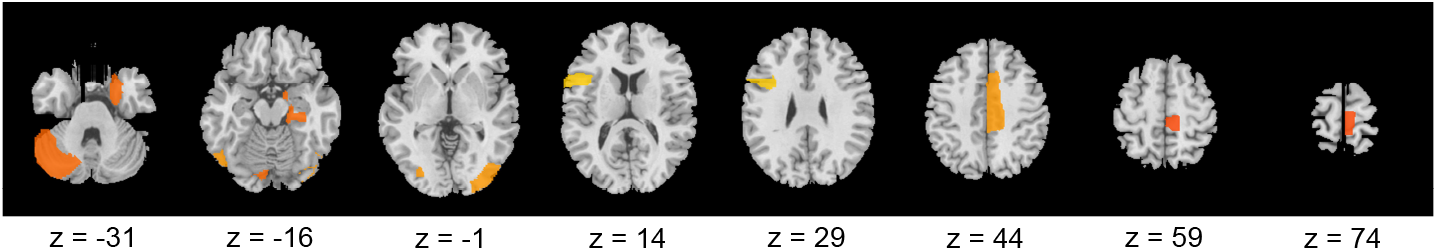
Classification brain regions between meditative state and resting state

## 4 Discussion

### 4.1 Performance comparison between T3Clus and the conventional approach

According to the conventional approach, t-test results revealed that there were no brain regions that significantly discriminated between resting and meditative states for all four feature values. This result suggests that it is difficult to explain the functional brain changes caused by meditation with a single functional brain metric.

On the other hand, when feature space was constructed considering the dependency between multiple variables by T3Clus, two states were classifiable and the discriminating features could be extracted. It is suggested that T3Clus can extract meaningful features that potentially affect classification between the two states. Additionally, these results also suggest that the short-term effects of breath-counting meditation on brain function in novice meditators are observed only in the projected space constructed by T3Clus.

### 4.2 Short-term effects of breath-counting meditation on brain functions

In Table 2, only the clustering coefficient was extracted as the discriminating feature. It quantifies the number of connections that exist between the nearest neighbors of a certain node as a proportion of the maximum number of possible connections [58]. With an increase in the clustering coefficient, the node tends to form with a high density of connections. In addition, it is notable that fALFF was not extracted. This suggests that local activity is not altered by the short-term effects of breath-counting meditation, while the functional network structure is affected.

**Table 2:** First component loading of feature values (*p* < 0.05)

| Feature value | loading | p-score |
| --- | --- | --- |
| Clustering coefficient | 0.7943 | 0.0370 |

In Table 1 and Table 2, all the component loadings of the eight brain regions and the feature value (clustering coefficient) constructing the first component were positive. Component scores can be calculated by the product of the observation values and the weights, which are obtained by the product of the loadings of the brain regions and the loading of the feature value. Therefore it indicates that the clustering coefficients of the eight brain regions are positively correlated with the first component score. Since the first component could discriminate between resting and meditative states, and the meditative states were distributed on the positive side of the first component in Figure 3, the clustering coefficients of the eight brain regions in the meditative state were higher than those in the resting state.

Next, we give an interpretation to each of the eight brain regions in Table 2. The right parahippocampal gyrus (ParaHippocampal R) located at the limbic system was extracted as one of the discriminating regions and plays an important role in creation of memories and recall of visual scenes [59]. It is also known as a part of the DMN [60]. Furthermore, it has been reported that the left crus I of the cerebellar hemisphere (Cerebellum Crus1 L) is a part of the DMN [61,62]. The inferior occipital cortex (Occipital Inf) is included in the visual regions of the occipital region [63–65] and also is a part of the DMN [66,67]. It has been reported that the DMN is activated in MW states, where the participant’s attention is distracted from breathing [20]. Our results suggested that the DMN regions, the ParaHippocampal R, Cerebellum Crus1 L, and Occipital Inf formed dense connections with other regions in moving from the resting state to the meditative state.

The right middle cingulate gyrus (Cingulum Mid R) is included in a fronto-parietal network (FPN), which is involved in top-down attention and controlling task execution [24, 25]. The opercular part of left inferior frontal gyrus (Frontal Inf Oper L) is also included in the FPN. It has been shown that the FPN is formed when the meditators sustain their attention on their breathing appropriately [20]. Therefore, it is expected that the densities of the networks that include the FPN regions, Cingulum Mid R, and Frontal Inf Oper L as nodes would be increased by sustaining the attention on breathing during meditation.

In addition, the right paracentral lobule (Paracentral Lobule R) is part of the somatosensory network (SSN) [68,69], and it has been reported that this network is activated when the meditator’s attention shifts back to the breath after being aware of MW [20]. This suggests that the Paracentral Lobule R is densely connected with other regions because the participants noticed MW in the meditation block and tried to return their attention to breathing again.

Each of three networks, DMN, FPN, and SSN, are known as characteristic functional network associated with the cognitive cycle that occurs during meditation. Based on our findings, we hypothesize that the short-term effect of breath-counting meditation is that each of the eight brain regions, mainly those included in the three networks, form a densely-connected network.

## 5 Conclusion

In this study, we investigated the short-term effects of focused-attention meditation on functional brain state in novice meditators. In the experiment, breath-counting meditation, one of the most popular forms of focused-attention meditation, was used, and brain activity during resting and meditation states was measured by fMRI. Functional changes in brain states were analyzed by the T3Clus method applied to the three graph theoretical metrics, degree centrality, betweenness centrality, and clustering coefficient and one spontaneous local activity measure, fALFF, calculated from the fMR images measured.

The results indicated that the two experimental conditions could be differentiated based on the first component of the two-dimensional feature space identified by T3Clus. Moreover, the component loadings of the first component revealed that the clustering coefficients of the eight regions were the prominent features to discriminate between resting and meditative states. We also found that the clustering coefficients of these regions tended to be higher in the meditation state. The extracted regions were included in either of three networks, DMN, FPN, and SSN, which are known to be characteristic functional network associated with the cognitive cycle that occurred during meditation. Our results revealed that the short-term effects of breath-counting meditation can be explained by the network density changes in these eight brain regions. It should be noted that no significant differences have been found, by t-test, in each of the four feature metrics for each brain region between two experimental conditions. This suggests that only T3Clus could extract the short-term effects by removing the feature metrics and the brain regions irrelevant to the functional changes caused by meditation.

## Ethics Statement

This study was carried out in accordance with the research ethics committee of Doshisha University, Kyoto, Japan (approval code: 15098) and was conducted in accordance with the Declaration of Helsinki.

## Conflict of Interest Statement

The authors declare that the research was conducted in the absence of any commercial or financial relationships that could be construed as a potential conflict of interest.

## Author Contributions

TM, TH and SH designed the study and wrote the manuscript. TM, KT, SY and SH conducted data analysis. HY and TH advised on the proposed method. All authors reviewed the manuscript.

## Funding

This work was supported by JSPS KAKENHI Grant Number JP19K12145.

