## Supplementary figures and images for "Short-term effects on brain functional network caused by focused-attention meditation revealed by Tucker3 clustering on graph theoretical metrics"

### Supplementary Figure S1

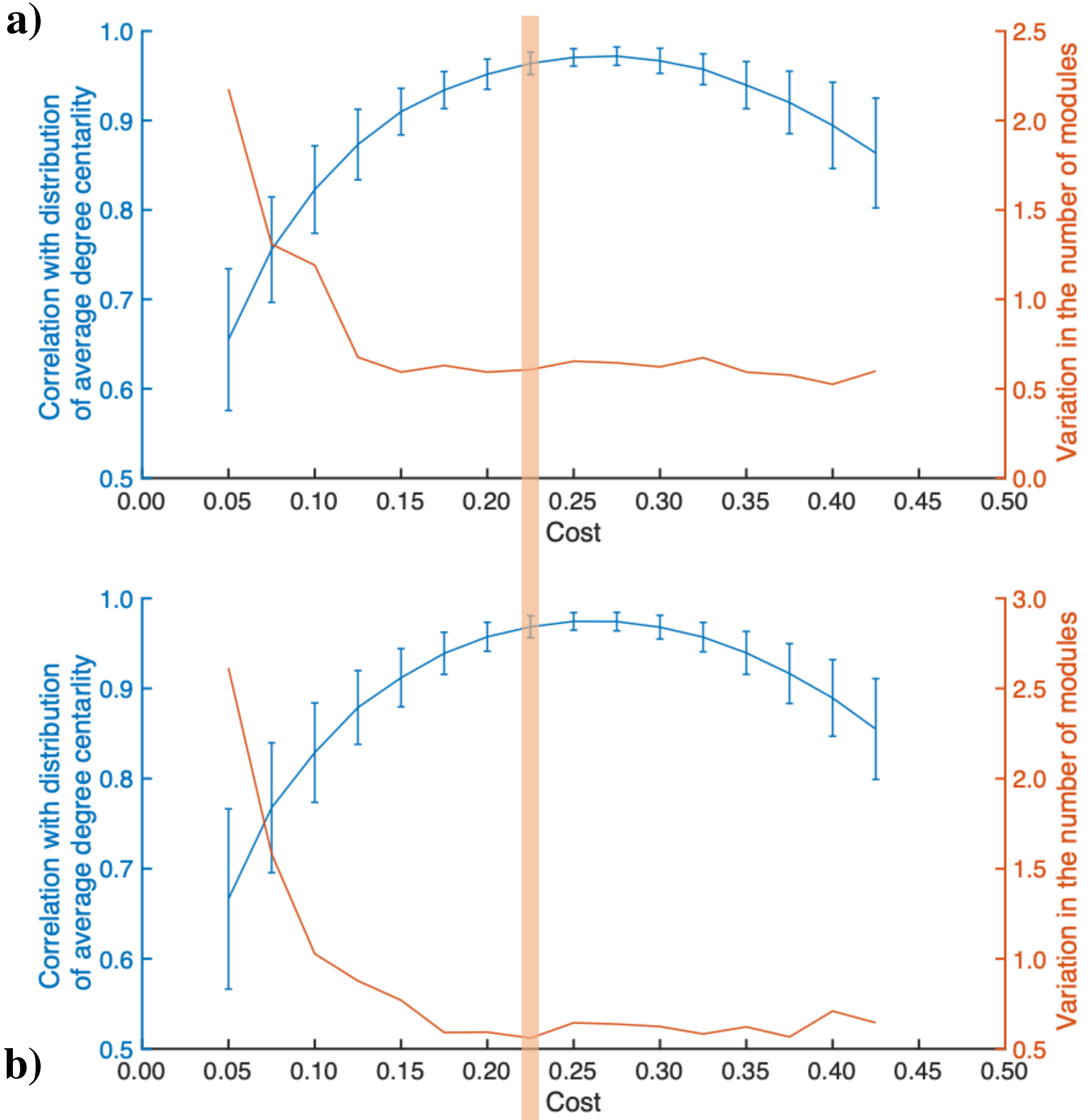
